# Influenza A virus within-host evolution and positive selection in a densely sampled household cohort over three seasons

**DOI:** 10.1101/2024.08.15.608152

**Authors:** Emily E. Bendall, Yuwei Zhu, William J. Fitzsimmons, Melissa Rolfes, Alexandra Mellis, Natasha Halasa, Emily T. Martin, Carlos G. Grijalva, H. Keipp Talbot, Adam S. Lauring

**Author notes:** Correspondence: Adam Lauring, University of Michigan, 1137 Catherine St., MS2 Room 4742C, Ann Arbor, MI 48109.

## Abstract

While influenza A virus (IAV) antigenic drift has been documented globally, in experimental animal infections, and in immunocompromised hosts, positive selection has generally not been detected in acute infections. This is likely due to challenges in distinguishing selected rare mutations from sequencing error, a reliance on cross-sectional sampling, and/or the lack of formal tests of selection for individual sites. Here, we sequenced IAV populations from 346 serial, daily nasal swabs from 143 individuals collected over three influenza seasons in a household cohort. Viruses were sequenced in duplicate, and intrahost single nucleotide variants (iSNV) were identified at a 0.5% frequency threshold. Within-host populations were subject to purifying selection with >75% mutations present at <2% frequency. Children (0-5 years) had marginally higher within-host evolutionary rates than adolescents (6-18 years) and adults (>18 years, 4.4x10^-6^ vs. 9.42x10^-7^ and 3.45x10^-6^, p <0.001). Forty-five iSNV had evidence of parallel evolution, but were not overrepresented in HA and NA. Several increased from minority to consensus level, with strong linkage among iSNV across segments. A Wright Fisher Approximate Bayesian Computational model identified positive selection at 23/256 loci (9%) in A(H3N2) specimens and 19/176 loci (11%) in A(H1N1)pdm09 specimens, and these were infrequently found in circulation. Overall, we found that within-host IAV populations were subject to purifying selection and genetic drift, with only subtle differences across seasons, subtypes, and age strata. Positive selection was rare and inconsistently detected.

## Introduction

Influenza A virus (IAV) evolution is dominated by antigenic drift. As the population gains immunity to circulating strain(s), antigenically distinct strains continue to emerge, displacing previous strains and forcing a change in the annual vaccine strain^1,2^. Accordingly, understanding the drivers of recurrent positive selection of antigenic variants is important for vaccine strain selection^3–5^. All genetic variation that is present in the virus population, including new antigenic variants, is ultimately derived from mutations that arise within individual hosts. Within host processes may at times parallel those seen at a global scale, as partial immunity from vaccination or previous infections may act as a selective force promoting the evolution of new antigenically distinct variants^6,7^.

As with many acute viral infections, the within-host dynamics of influenza virus are dominated by purifying selection and characterized by low levels of genetic diversity^8–14^. However, there is mixed evidence for specific sites being under positive selection. Several studies have reported nonsynonymous mutations in antigenic sites^8–12^, and these sites are occasionally shared among individuals^9,10^, exist at high frequency^9,11,12^, or are found at detectable levels in the global population^8,12^. More often, these mutations are identified only in single individuals at low (<5%) frequencies. Additionally, the frequency and number of mutations tend not to differ between antigenic and non-antigenic sites^8,13^.

Within-host selection may not be uniform across host populations. While we have found that vaccination has minimal impact on patterns of within-host diversity, other factors, such as age, have not been thoroughly explored^10^. Children have prolonged shedding compared to adults, and the extra time may allow for mutations in antigenic sites and their subsequent selection^15^. A recent study focused on young children (<5 years old), took place in Southeast Asia where child vaccination rates are very low, and most children didn’t have detectable antibody titers in the study^11^. Mutations were identified in antigenic sites, but there wasn’t a clear signal of positive selection. By contrast, most adolescents will have had multiple exposures to influenza^16^, with a shedding period that is of intermediate length^15^. This combination may be the optimal for the selection of new antigenic variants within hosts, but there haven’t been any studies focused on this group.

There are several technical reasons why positive selection has been difficult to detect. Selection is difficult to detect and quantify with cross-sectional sampling, and most studies have reported 1 or 2 samples per subject without longitudinal sampling^12^. Second, the frequency threshold for detecting intrahost single nucleotide variants (iSNV) is typically 2%. Most intra-host variants are *de novo* mutations due to tight bottlenecks during transmission and the detection threshold may be too high to detect many of these iSNV^8,14,18^. Finally, there are few formal tests for positive selection on a per site basis; most interrogate on a gene level (e.g., nucleotide diversity), or informally on a per site basis (e.g., iSNV at important sites). In some cases, single mutations are sufficient to cause antigenic drift making per site tests necessary to detect selection for antibody escape^19^.

Here we address the design and technical limitations of previous studies. We developed and benchmarked a sequencing and bioinformatic pipeline that allowed us to accurately identify iSNV at a frequency threshold of 0.005 (0.5%). We applied this method to individuals sampled within a case-ascertained household cohort, in which nasal swab specimens were collected every day for 7 days after symptom onset. The study covered three influenza seasons and included individuals across the age spectrum with a mixture of antiviral treatment and vaccination histories. We use these sequence and host data to define patterns of within-host genetic diversity and divergence with respect to age, vaccination, antiviral usage, and timing of sample collection. We applied multiple methods to detect selection on a per site basis with clear statistical cutoffs.

## Methods

### Cohort and Specimens

Households were enrolled through the Influenza Transmission Evaluation Study (FluTES), a case ascertained household transmission study based in Nashville, TN that enrolled over the 2017/18, 2018/19, and 2019/20 Northern Hemisphere influenza seasons^20^. All individuals provided informed consent, and the study was approved by the Vanderbilt University Medical Center Institutional Review Board. The first household members with laboratory-confirmed IAV infection (index cases) were identified and recruited from ambulatory clinics, emergency departments, or other settings that performed influenza testing. For this study, we focused on IAV only. Index cases with acute illness of <5 days duration who lived with at least 1 other person who was not currently ill were eligible to participate. The index case and their household contacts were enrolled within 7 days of the index case’s illness onset. Influenza vaccination was self-reported at enrollment and was included if both date and location of vaccination were provided. Participants who reported vaccination within the prior 14 days had their vaccine status listed as unknown. For individuals with an asymptomatic infection, the symptom onset date of the index case was used as that individual’s onset date. Individuals were divided into three age groups: children (≤5 years), adolescents (6-17 years), and adults (≥18 years) for further analyses.

Nasal swabs were self-/parent-collected daily during follow-up for 7 days and tested for influenza using reverse transcriptase quantitative polymerase chain reaction (RT-qPCR) at Vanderbilt University Medical Center using the CDC Human Influenza Virus Real-Time RT-PCR Diagnostic Panel, Influenza A/B Typing Kit with the SuperScript III Platinum One-Step qRT-PCR Kit (Invitrogen) on the StepOnePlus or QuantStudio 6 Flex (Applied Biosystems). Subtyping of IAV-positive specimens was performed using the CDC Human Influenza Virus Real-Time RT-PCR Diagnostic Panel, Influenza A Subtyping Kit with the SuperScript III Platinum One-Step qRT-PCR Kit System on the MagNA Pure LC 2.0 platform (Roche).

### Benchmarking variant calling

We used data from McCrone et al. 2016 to benchmark our variant calling pipeline^21^. Briefly, 20 viruses, each with a single point mutation, were generated in a WSN33 background. The 20 mutant viruses were mixed with wild type WSN33 to create populations in which each mutant was present at 5, 2, 1, 0.5%. These samples were processed and sequenced in duplicate on an Illumina Miseq. We aligned the data to WSN33 using Bowtie2^22^ with the “very sensitive” setting and duplicate reads were discarded using picard tools^23^. Reads from both replicates were combined and used to make a within host consensus sequence using a script from Xue et al.^13^. The replicates were separately aligned to this consensus and duplicates were removed. iSNV were called using iVAR in each replicate^24^. To be considered for variant calling, reads had to have a mapping quality ≥20, and bases had to have a phred score ≥30. iSNV had to have a per-site sequencing depth of ≥400, and an iVAR p-value ≤1x10^-5^. iSNV were retained only if they were found in both sequencing replicates. We calculated the specificity and sensitivity for iSNV detection at each frequency threshold (0.5%, 1%, 2%, 5%).

### Sequencing

IAV-positive samples with an RT-qPCR cycle threshold (Ct) value ≤30 were sequenced in duplicate after the RNA extraction step. RNA was extracted using Invitrogen PureLink Pro 96 Viral RNA/DNA Purification Kits on an EpiMotion or a MagMAX viral/pathogen nucleic acid purification kit (ThermoFisher) on a Kingfisher Machine. SuperScript IV one-step RT-PCR kits and universal IAV primers were used for RT-PCR^25^. Library preparation was completed by using the Illumina DNA prep kit, and libraries were sequenced on a Novaseq (2x150 PE reads) by the Advanced Genomics Core at the University of Michigan.

Reads from each sample were aligned to the vaccine strain for each subtype and year for the initial alignment. For 2019/20 A(H1N1)pdm09 we used A/New Jersey/13/2018, since the A/Brisbane/02/2018 sequence was not available. Variant calling was performed as above on samples in which each replicate had an average genome-wide coverage of at least 1000x with an iSNV frequency cut-off of 0.005 (0.5%). For iSNV in overlapping ORFs, an iSNV was classified as a nonsynonymous if it was nonsynonymous in any ORF. Stop codons were classified as nonsynonymous. For all further analyses we used the average iSNV frequency in the two replicates as the iSNV frequency.

### iSNV Dynamics and Divergence rates

We calculated the divergence rate using the methods from Xue and Bloom 2020^13^. Briefly, we calculated the rate of evolution by summing the frequencies of within-host mutations and divided by the number of available sites and time since the infection began. To account for iSNV that go from minor to major allele in individuals with multiple samples, the allele frequency used was from the allele that was minor in the earliest sample. We calculated the rates separately for nonsynonymous and synonymous mutations. We used 0.75 for the proportion of available sites for nonsynonymous mutations and 0.25 for synonymous. To determine the number of available sites, we multiplied the proportion of sites available by the length of the coding sequence for the relevant reference. We excluded M and NS segments, because overlapping open reading frames allow an individual mutation to be both synonymous and nonsynonymous. Because symptoms typically start 2-3 days post infection, we added 2 days to the time since symptom onset to get the time since infection began^26–28^. We excluded individuals who were asymptomatic from the divergence rate analysis. We also excluded outlier samples that had ≥50 iSNV.

We calculated the rate of evolution for each sample with nonsynonymous and synonymous rates calculated separately. Because the calculated rate of divergence varied over the course of the infection, we also calculated the rate using the sample with the lowest Ct value for each individual. The rate was calculated for the whole genome and for each segment.

### Analysis of shared iSNV

We performed permutation simulations for each reference strain to determine the expected number of individuals who would share an iSNV based on the number of individuals, the number of iSNV, and the genome size^29^. We set the proportion of the genome that was mutable as 0.6 based on experimental data^30^. One thousand permutations were performed for each reference strain. For a given group (e.g. mutations shared between 2 individuals), we calculated the p value as the proportion of permutations that had as many or more shared iSNV than the observed number of shared iSNV. Because we were interested in the number of times that an iSNV independently arose, iSNV that were shared among multiple individuals within the same household were recorded only once for that household for both the simulations and the observed number of individuals. Samples and individuals with >50 iSNV were excluded.

### Wright Fisher Approximate Bayesian Computational method

We mapped the allele trajectory of alleles that changed from a minor to a major allele over the course of an infection. We also used Wright Fisher Approximate Bayesian Computation (WFABC) to estimate the effective populations size (Ne) and per locus selection coefficient (s) based on allele trajectories^31^. A generation time of 6 hours was used^26^. To maximize the number of loci used in the calculation of Ne and to avoid violating the assumption that most loci are neutral, we estimated a single Ne for A(H1N1)pdm09 and for A(H3N2). We used all loci in which the first two time points were one day apart to estimate Ne. 10,000 bootstrap replicates were performed.

A fixed Ne was used for the per locus selection coefficient simulations, with the analysis repeated for the mean Ne, and +/- 1 standard deviation estimated from the previous step. A uniform prior between s of -0.5 and 0.5 was used. 100,000 simulations with an acceptance rate of 0.01 was used. We estimated the 95% highest posterior density intervals using the boa package^32^ in R. We considered a site to be positively selected if the 95% highest posterior density didn’t include 0 for all three effective population sizes.

### Statistical Methods

Descriptive statistics were conducted. The Mann-Whitney U tests were used to compare the number iSNV per sample and iSNV frequencies by mutation type, vaccination, and antiviral usage. The Kruskal-Wallace tests were performed for age and days post symptom onset. For divergence rates, Mann-Whitney U tests were performed for vaccination, antiviral usage, and IAV subtype. Kruskal-Wallace tests were performed for age, segment, and days post infection. We also compared the likelihood of a shared iSNV or a positive selection coefficient. χ^2^ tests were performed to test if age, vaccination, and antiviral usage affected the likelihood of an individual having a shared iSNV or impacted the probability of an individual having at least 1 iSNV with a positive selection coefficient.

All analyses were conducted using R version v4.2.0.

### Data and Code Availability

Raw sequence data are available at the NCBI Sequence Read Archive (SRA) under Bioproject PRJNA1085292. Analysis code is available at https://github.com/lauringlab/Flutes_within-host_evolution

## Results

### Benchmarking

The FluTES study was a case-ascertained household cohort that enrolled over the 2017/18 through 2019/20 Northern Hemisphere influenza seasons. Consistent with the viruses circulating in the United States during this timeframe, the 2017/18^33^ and 2018/19^34^ seasons were A(H3N2) predominant and the 2019/20 flu season was exclusively A(H1N1)pdm09^35^. In total, there were 302 cases at the Vanderbilt site over these three seasons, and the majority provided 7 daily specimens. We successfully sequenced 346/413 (84%) specimens from 143 individuals (Table 1). Eighty-three out of 143 (58%) individuals had multiple sequenced specimens (Figure S1). Among the 143 individuals, there were 37 children (≤5 years), 51 adolescents (6-17 years), and 55 adults (≥18 years).

**Table 1.** Overview of specimen types.

| Subtype | Season | Number of Specimens | Number of Individuals |
| --- | --- | --- | --- |
| **H3N2** | 2017-2018 | 32 | 13 |
|  | 2018-2019 | 153 | 58 |
|  | 2019-2020 | 0 | 0 |
| **H1N1** | 2017-2018 | 13 | 7 |
|  | 2018-2019 | 35 | 14 |
|  | 2019-2020 | 113 | 51 |

In order to accurately detect ultra-rare mutations, we refined our variant calling approach and benchmarked this new method against a previously described dataset^21^. In our benchmarking, we found high specificity across all mutant frequencies, but sensitivity decreased as mutant frequency decreased (Table 2). To detect the greatest number of true variants, we used a 0.005 frequency threshold for the rest of the study. We sequenced each specimen in duplicate, obtaining high coverage across the genome (mean >10,000x) and consistency in iSNV frequency between the sequencing replicates (Figure 1A,B). iSNV were more common in the 3^rd^ codon position compared to the 1^st^ and 2^nd^ position at frequencies above our 0.5% cutoff. As sequencing errors are presumably randomly dispersed, this further suggests that we are detecting true iSNV (Figure 1C).

**Figure 1.**
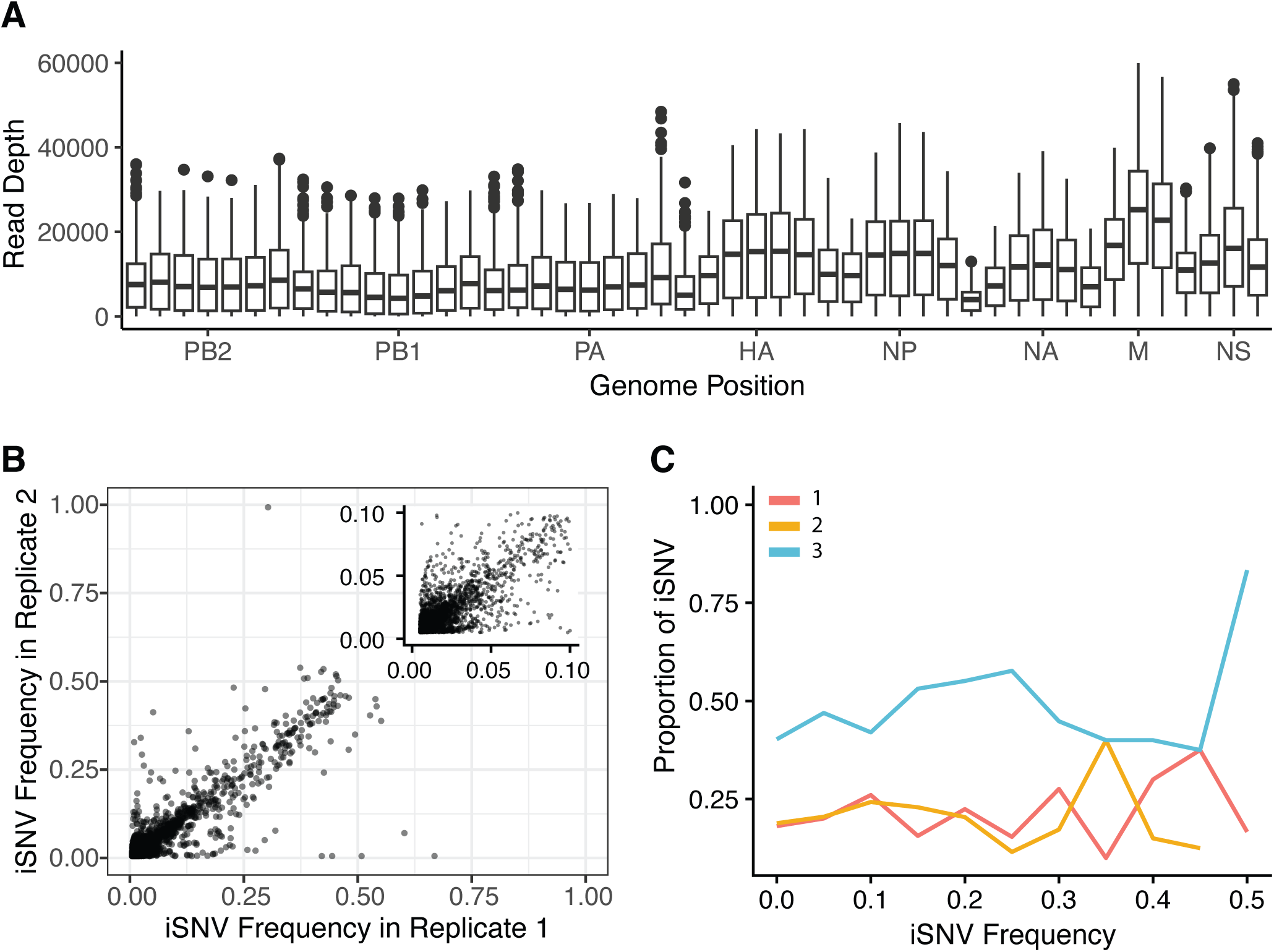
**(A).** Sequencing coverage across the genome in 300 bp nonoverlapping windows. **(B).** iSNV frequency is consistent across replicates. The insert shows iSNV frequency up to 0.1. **(C).** Proportion of iSNV in each codon position for a given frequency in 0.05 frequency bins. iSNV in codon position 3 (blue) are more common than iSNV in position 1 (red) or 2 (gold) across the frequency range.

**Table 2.** Benchmarking results.

| % Freq | True<br>positives | Sensitivity | False<br>Positives | Specificity |
| --- | --- | --- | --- | --- |
| 5 | 18 | 0.9 | 13 | 0.9996759 |
| 2 | 15 | 0.75 | 6 | 0.9998504 |
| 1 | 13 | 0.65 | 10 | 0.9997507 |
| 0.5 | 6 | 0.3 | 2 | 0.9999501 |

### Cross sectional analysis

Most specimens had between 1-50 iSNV at >0.5% frequency, while 4 specimens had 0 iSNV and 5 specimens had >50 iSNV (Figure 2A, Table S1). Children had more iSNV per specimen (median 11) than adolescents (median 7, p=0.001, Figure S2). The number of iSNV per specimen increased as the infection progressed before decreasing again, mirroring typical viral titer trajectories (Figure S2). However, the difference in the number of iSNV wasn’t significant. The number of iSNV didn’t vary much by vaccine status or antiviral usage (Figure S2).

**Figure 2.**
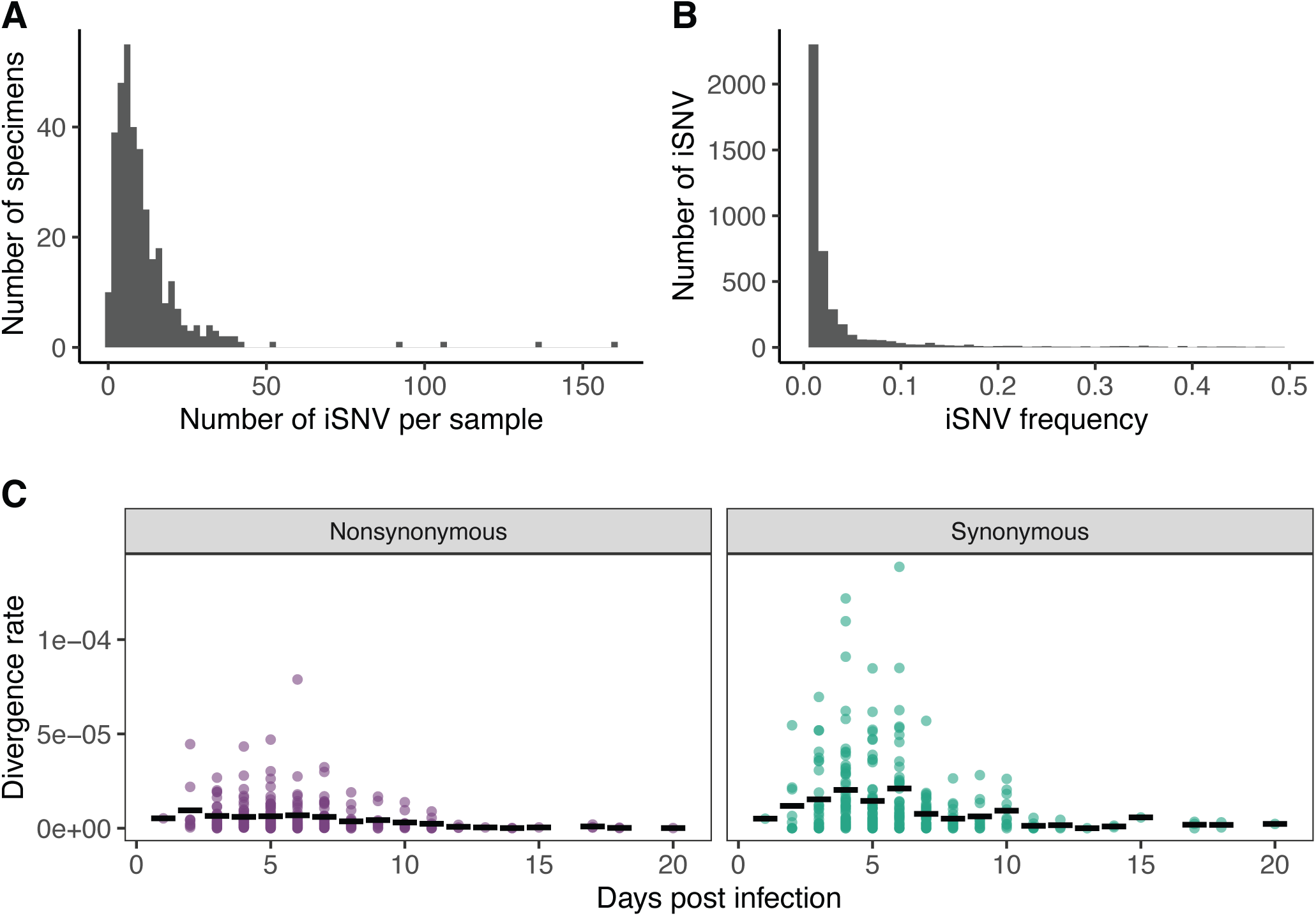
**(A).** Number of iSNV per specimen. **(B).** iSNV frequency**(C).** Divergence rate (divergence/site/day) by day post infection (green synonymous, purple nonsynonymous).

The vast majority of iSNV tended to be very low frequency and most were between 0.5 and 2% (Figure 2B). The frequency of iSNV varied significantly by day of sampling post symptom onset (p<0.001), host age (p<0.001), mutation type (p=0.005), vaccine status (p<0.001) and antiviral usage (p=0.013, Table S1, Figure S3). Adults had the lowest median frequency (0.013), followed by children (0.014) and adolescents (0.015, Figure S3). Nonsynonymous iSNVs (median 0.015) were found at slightly higher frequencies than synonymous (median 0.013, Figure S3). Specimens from vaccinated individuals (median 0.015) had higher median iSNV frequencies than unvaccinated individuals (median 0.013, Figure S3C), while specimens from those receiving antivirals had lower iSNV frequencies (median 0.013 vs 0.014, Figure S3B). While all of these differences were statistically significant, the magnitude of the differences were extremely small, and the iSNV frequency distributions almost completely overlapped.

We estimated within-host evolutionary rates as nucleotide divergence per site per day^13^. Consistent with the expansion and contraction of the viral population over the course of an infection, the estimated divergence rate varied according to the day of sampling (Figure 2C, Table S2). The divergence rate increased from the onset of the infection until approximately day 5 and then decreased. This time-varying pattern was more pronounced for synonymous than nonsynonymous mutations. Children (0-5 years) had marginally higher within-host evolutionary rates than adolescents (6-18 years) and adults (>18 years, 4.4x10^-6^ vs. 9.42x10^-7^ and 3.45x10^-6^, p <0.001). (Figure S4). A(H1N1)pdm09 infections had a faster divergence rate than A(H3N2) for nonsynonymous mutations (p= 0.010, Figure 2). The divergence rate did not vary by genome segment (Figure 2), host antiviral usage, or host vaccination status (Table S2).

### Shared iSNV and Evolutionary Convergence

Evolutionary convergence is a signal of positive selection and can be identified based on the sharing of iSNV among individuals who are not linked by transmission. In our analysis, we first determined the expected number of iSNV that would be shared based on chance alone given the number of sequenced specimens and iSNV for each subtype and season. The corresponding threshold for statistical significance was ≥2 shared iSNV for 2017/18 A(H3N2) and 2017/18 A(H1N1)pdm09, ≥4 for 2019/20 A(H1N1)pdm09, and ≥3 for 2018/19 A(H3N2) (p<0.001 for all reference strains, Figure S5). Nearly all shared iSNV were found at very low frequencies (Figure S6).

The hemagglutinin (HA) and neuraminidase (NA) genes did not exhibit an overabundance of shared iSNVs, and the frequencies of shared iSNVs in HA and NA were not higher than iSNV in other genes. Only one shared iSNV was within an antigenic site, and it was synonymous (T620C [G197]). There was an overrepresentation of shared iSNV in 2017/18 A(H3N2) M and NS with 28% and 45% of shared iSNV despite these segments making up 8% and 6% of the genome respectively. There was also an overrepresentation in 2017/18 & 2018/19 A(H1N1)pdm09 PB1 and M. They have 37% and 26% of the iSNV, but only make up 17% and 8% of the genome respectively.

High iSNV specimens (>50 iSNV) were responsible for 156/201 (78%) of shared iSNV. Because these very high numbers of iSNV are inconsistent with the typical rate of mutation accumulation and are plausibly due to low-level (<1%) contamination or coinfection, we repeated this analysis after removing specimens with >50 iSNV. There was no effect on the number of shared iSNV in HA and NA (Figure 3). The number of iSNV in 2017/18 & 2018/19 A(H1N1)pdm09 PB1, M, and 2017 A(H3N2) M segments were greatly decreased, but the overrepresentation of shared iSNV in the 2017 A(H3N2) NS segment remained. When we plotted shared iSNV in NS by individual, we found that most of the shared iSNV were part of a low frequency, ∼400 bp haplotype shared among 9 individuals (Figure S6).

**Figure 3.**
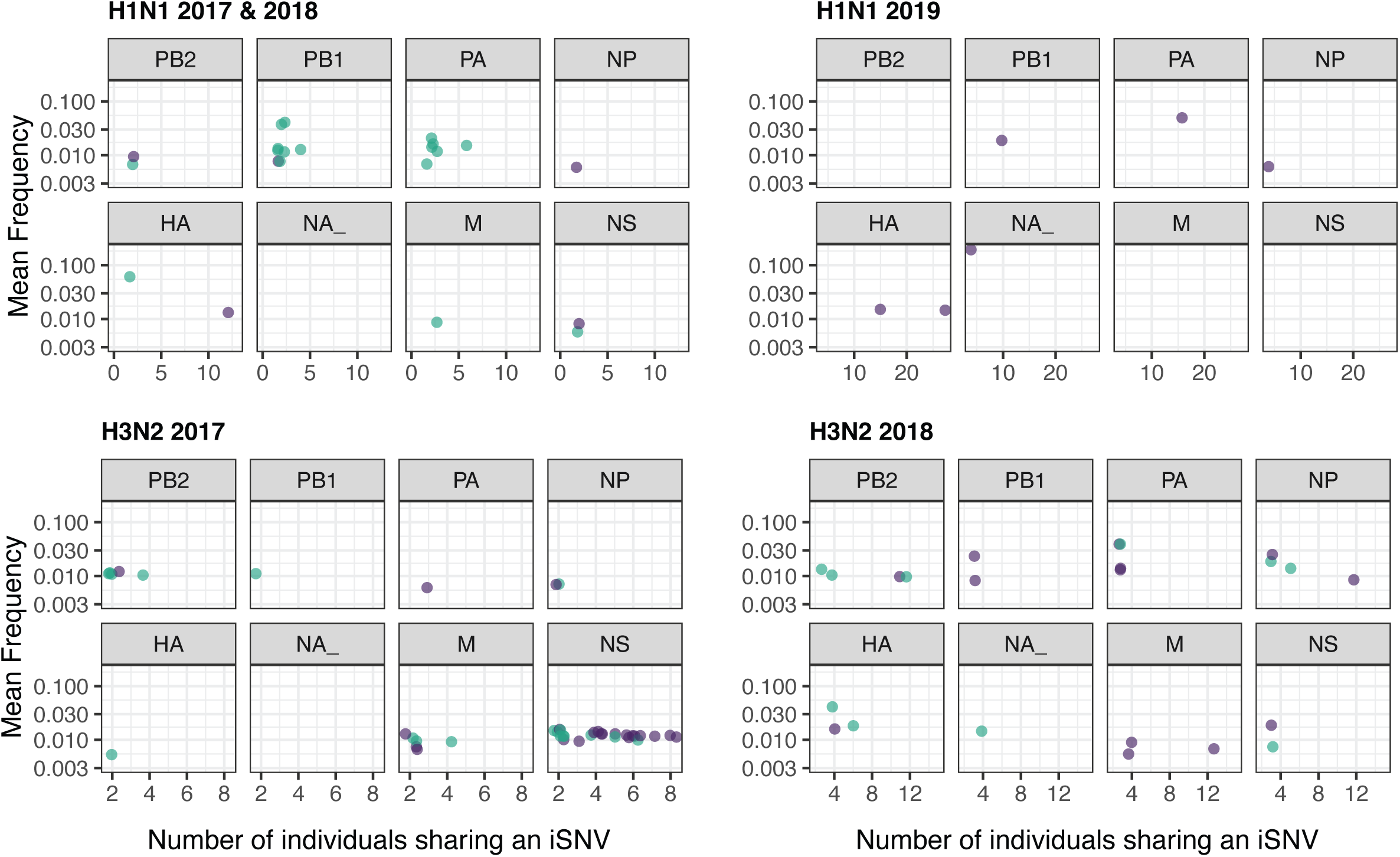
The mean frequency of iSNV found in multiple individuals from different households after samples with >50 iSNV were removed. iSNV from 2017/18 & 2018/19 H1N1 and 2017/18 H3N2 infections are shared in ≥ 2 individuals. iSNV from 2019/20 H1N1 infections are shared in ≥ 3 individuals, and iSNV from 2018/19 H3N2 infections are shared in ≥ 4 individuals. (Green synonymous iSNV, purple nonsynonymous iSNV)

Vaccination and antiviral usage did not affect the likelihood of an individual contributing to a shared variant (Table S3). Host age did affect the likelihood, but no pairwise comparisons achieved statistical significance; 17% of iSNV in adults were shared, 14% in adolescents, and 12% in children.

### Analysis of selection in serial specimens

Thirty iSNV changed from a minor to major allele during an infection. This included 2 synonymous (G665A [A212] and G686A [T219]) and 2 nonsynonymous iSNV (C556A [T176K] and G564A [A179T]) in HA antigenic sites from A(H3N2) infections. Importantly, there was evidence for linkage disequilibrium and hitchhiking, with neutral synonymous iSNV being swept along with putatively selected nonsynonymous iSNV^36^. Eight individuals had >1 iSNV that became the major allele, and in many cases the allele trajectories of these iSNV closely matched each other, even when present on different segments (Figure 4A). For example, individual 1811602 (H3N2) had iSNV at 3 loci on HA, NP, and PB2 (◆ in Figure 4A) with very similar trajectories. The iSNV on HA and NP are nonsynonymous while the iSNV on PB2 is synonymous.

**Figure 4.**
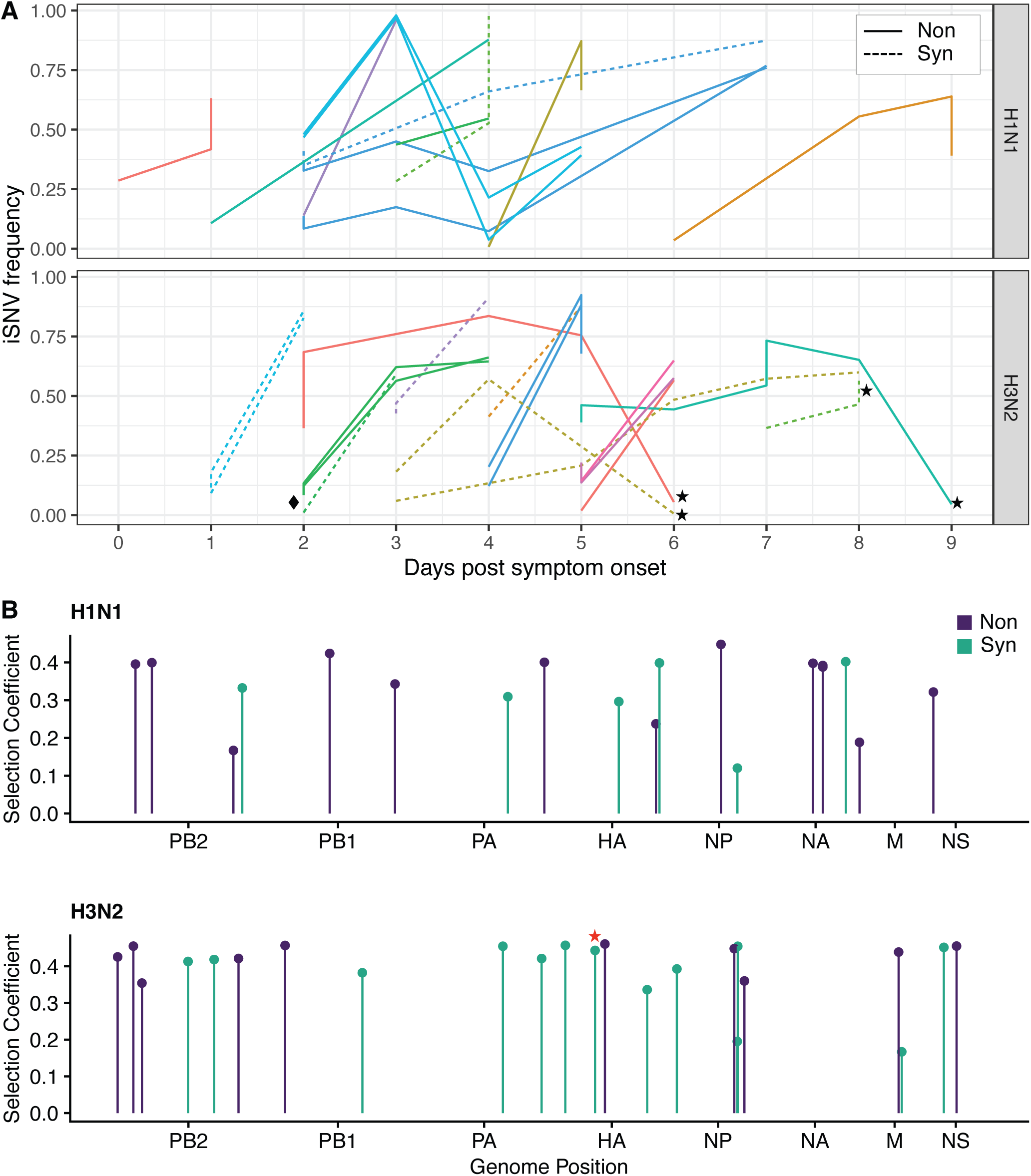
**A**. Allele trajectories of iSNV that go from minor to major allele over the course of an infection. Lines are colored by individual. Dashed lines are synonymous iSNV and solid lines are nonsynonymous iSNV. The household symptom onset date was used for individuals with asymptomatic infections. Green lines next to the diamond (◆) are from individual 1811602 **(B).** WFABC selection coefficients for iSNV under positive selection. Stars (!) indicate iSNV in HA antigenic sites in both **(A)** and **(B)**.

We used a Wright Fisher Approximate Bayesian Computational Method to infer selection coefficients on individual iSNV. For A(H1N1)pdm09 and H3N3, 129 and 199 loci were used to estimate the within-host effective population size, respectively. The inferred effective population size for A(H3N2) infections was 284 ± 60 and was 176 ± 41 for A(H1N1)pdm09 infections. Twenty-three out of 256 loci (9%) in A(H3N2) specimens and 19/176 loci (11%) in A(H1N1)pdm09 specimens had positive selection coefficients (Figure 4B). One A(H3N2) synonymous iSNV (T620C [G197]) was in an HA antigenic site and six A(H1N1)pdm09 and 13 A(H3N2) iSNV were synonymous. Three of these synonymous A(H1N1)pdm09 iSNV had corresponding nonsynonymous iSNV under positive selection. Thirteen A(H3N2) and 10 A(H1N1)pdm09 iSNV with positive selection coefficients also reached a majority allele frequency. Host age, vaccination status, and antiviral usage did not affect the likelihood of an individual having an iSNV with a positive selection coefficient (Table S4).

### Global trajectory

We next used the seasonal influenza Nextstrain builds to visualize whether any of the iSNV in HA and NA that are putatively under positive selection were also identified as increasing in global frequency^37^. Most of the iSNV identified in our study were either already a dominant allele in the season under evaluation or didn’t exceed 5% frequency at any timepoint. However, there were 2 A(H1N1)pdm09 iSNV and 4 A(H3N2) iSNV that circulated at a significant global frequency after appearing in our cohort (Figure S7). None of these were in antigenic sites. For A(H1N1)pdm09, both iSNV were from the 2019/20 season; HA_G1628A (A532) is a synonymous iSNV that had a positive selection coefficient. It had a global frequency of ∼4% in the 2019/20 season and was fixed in the global population by 2023. HA_A754C (E241A) is a nonsynonymous iSNV that was shared among 19 individuals from 16 households. It was not found at a detectable frequency globally until 2021 and was predominant in 2023. For A(H3N2), HA_T314C (F95) was found in 2 people during the 2017/18 season and HA_G225A was found in 4 people during the 2018/19 season. HA_T314C is synonymous and reached a frequency of 3% in 2017. It never became a dominant allele but did reach a frequency of 10% in 2021. It was previously the dominant allele during 2012 before HA_T314 became dominant. HA_G225A (E66K) is nonsynonymous and was not circulating during 2018. In 2019 it appeared and fluctuated between 0% and 25% until 2022, when it became the dominant allele. It reached 100% in 2024.

HA_C556A (T176K) and HA_G665A (A212) were minor alleles that became major alleles. HA_C556A is a nonsynonymous mutation that was found in the 2017/18 cohort. HA_G665A is a synonymous mutation found in the 2018/19 cohort. A556 fluctuated in frequency as a minor allele in the global population with C556 being the dominant allele, until both were replaced by T starting in 2020. The A allele has fluctuated in global frequency since our sample and has reached a maximum frequency of 20%. The G allele was not circulating at detectable levels when the sample was taken or in the subsequent year. The G was detected in 2023 at a maximum frequency of 2%.

## Discussion

In our detailed analysis of serially sampled individuals in a longitudinal household transmission study, we found that within-host IAV populations are highly dynamic and subject to strong purifying selection and genetic drift. There were differences in divergence rates and number of iSNV per sample based on age, and multiple factors influenced iSNV frequencies. However, these differences were minimal and weren’t reflected in differences in their selection. Positive selection was rare and inconsistently detected with the three methods applied. Our comprehensive evaluation demonstrates that in most cases, the extent of within-host evolution is small and positive selection is only a minor contributor.

Consistent with strong purifying selection, we found low genetic diversity and divergence rates, despite influenza virus’s high mutation rate. With the lower frequency threshold, we observed more iSNV than other studies, but they mainly occurred in the 0.005-0.02 frequency range. Our divergence rates were lower than the divergences rates calculated by Xue et al.^13^ While the same frequency threshold was used, we applied more rigorous variant calling and potentially had a different distribution of sample timing. Our divergence rates were similar to the rates reported by Han et al. 2021^11^.

By lowering the frequency threshold and using serial sampling we were able to detect iSNV that were putatively under selection. However, these iSNV were mostly outside of HA and NA, and very few were in known antigenic sites. Even when there were selected iSNV in HA and NA, they rarely achieved a significant frequency in the global population. The small effective population size of within-host populations reduces the efficacy of selection, and partial immunity may be a relatively weak selective force. If a virus escapes the mucosal membrane to successfully infect a host cell, it takes several days for an antibody mediated recall response^38^. The asynchrony of the infection and antibody response would result in minimal selection on antigenic sites within hosts initially. Here, the time between the antibody response and viral clearance may not be sufficient for significant selection. Weak selection combined with a short duration of infection in acutely infected individuals minimizes the evolution of new antigenic variants. We do not deny that new antigenic variants can arise through within-host selection. However, they will be rare events, and newly arising variants, even in antigenic sites, can sweep to fixation due to stochastic dynamics.

Our study focused on typical influenza virus infections from seasonal viruses, but the evolutionary dynamics may differ in atypical situations where positive selection may be prominent. During prolonged infections in some immunocompromised individuals, an antibody mediated recall response is mounted before the virus is cleared. Here, within-host antigenic selection parallels population-level selection and mutations that arise within hosts are seen at high frequency globally in subsequent seasons^39^. Similarly, when positive selection is strong, an acute infection is long enough for adaptive mutations to arise and increase in frequency. Antiviral mutations associated with oseltamivir resistance are routinely seen in acute infections during treatment^11,12,40,41^. Additionally, during the first wave of the 2009 A(H1N1)pdm09 pandemic, there was an overabundance of nonsynonymous mutations within hosts^11^. As a new zoonotic spill over, the virus was not already adapted to human hosts allowing for stronger selection^42^. Based on our prior work and those of others, we suspect that the same principles will generalize for most self-limited respiratory virus infections.

Because our study spanned three influenza seasons and was significantly larger than prior ones, we were able to evaluate the impact of viral and host factors on within-host diversity and evolutionary rate. Consistent with the observed differences in shedding, we observed differences in the levels of genetic diversity across age groups. Children had faster rates of evolution and a greater number of iSNV per specimen compared to adolescents, while adolescents had higher iSNV frequencies than adults or children. Although these comparisons achieved statistical significance, they are unlikely to be biologically meaningful. The differences between ages are very small and had minimal impact on our tests for selection. In our cohort, positively selected iSNV were equally likely to have been identified in children, adolescents or adults; children were less likely to contribute to shared iSNV than adults. Although children and adolescents may play an outsized role in influenza epidemics due to increased shedding and their social networks^43^, their within-host dynamics are remarkably similar to those of adults.

Our study has multiple strengths. The participants were from the community and were representative of the general population, making the results widely applicable. Due to the diversity in our cohort, we were able to study the effects of age, vaccination status, and antiviral usage. We also used multiple methods to detect positive selection. Allele trajectory methods are more effective, but identifying shared variants allowed us to test for positive selection when there was only a single specimen. Additionally, we had applied stringent variant calling criteria that were benchmarked to ensure high confidence in variant calls.

There are several limitations to our study. We used only nasal swabs and would potentially miss selected sites if there was compartmentalization. However, nasal swabs likely sample the most relevant population for transmission, as virions replicating in the soft palate or nasal epithelial cells form the population that is most likely to be transmitted^44–46^. In ferrets, compartmentalization in the lungs occurs through a series of bottlenecks^47^. This leads to significant genetic drift, which further masks any signals of positive selection. Therefore, it is unlikely that more comprehensive sampling of the respiratory tract would change our results. Despite our best attempts to mitigate it, there is always the possibility of sequencing error and false iSNV calls.

Our work was only in a US cohort, and there is the possibility that the results will not generalize to other settings. However, the viruses circulating in our cohort were representative of seasonal influenza viruses globally^48–50^ and many cases were not medically attended. Additionally, in 2020, the widespread mitigation measures in response to the COVID-19 pandemic severely curtailed influenza A virus circulation^51^. Influenza A virus reappeared in 2021 after an intense bottleneck. In contrast to typical patterns, stochastic processes dominated globally, complicating our ability to connect within-host processes to global outcomes. However, our results are similar to other acute seasons and all A(H3N2) specimens were collected 1 to 2 seasons prior to the SARS-CoV-2 outbreak.

Despite multiple studies with varying populations and approaches, clear evidence of within-host antigenic drift in acute infections is lacking. Given the rigor of our methods and the data from other studies, we find that acute influenza virus infections are dominated by stochastic forces (mutation and drift) with little evidence for positive selection regardless of age, vaccination status, or antiviral use, or subtype. This calls into question whether this is useful as a surveillance strategy. While positively selected variants exist, a large number of people would need to be closely followed to find them. However, our characterization of divergence rates within-hosts as well as the impact of viral and host factors is important for understanding and modeling how within-host processes feed into larger evolutionary dynamics. Other RNA respiratory viruses, such as Influenza B^29^, SARS-CoV-2^52–56^ and RSV^57^, have low within-host genetic diversity and similar infection dynamics to IAV. The phenomenon of strong immune selection at the population level and stochastic processes and purifying selection dominating within hosts may be widespread in acute respiratory infections.

## Funding Acknowledgement

Primary funding for the FluTES study was provided by the US Centers for Disease Control and Prevention (CDC 5U01IP001083). CGG was partially supported by NIH K24A I148459. Scientists from the US CDC participated in all aspects of this study, including its design, analysis, interpretation of data, writing the report, and the decision to submit the article for publication. Sequencing and associated analysis was supported by a Burroughs Wellcome Fund Investigator in the Pathogenesis of Infectious Diseases Award (to ASL), NIH R01 AI148371 (to ASL and ETM), and the Penn Center for Excellence in Influenza Research and Response, Penn-CEIRR, NIH 75N93021C00015 (to ASL and ETM).

## Disclaimer

The findings and conclusions in this report are those of the authors and do not necessarily represent the official position of the Centers for Disease Control and Prevention (CDC).

## Author Contributions

Conceptualisation: Lauring, Grijalva, Talbot

Data Curation: All authors

Formal Analysis: Bendall, Zhu, Lauring

Funding Acquisition: Lauring, Martin, Grijalva, Talbot

Investigation: All authors

Methodology: All authors

Project Administration: Lauring, Grijalva, Talbot

Resources: All authors

---

**Figure S1.**
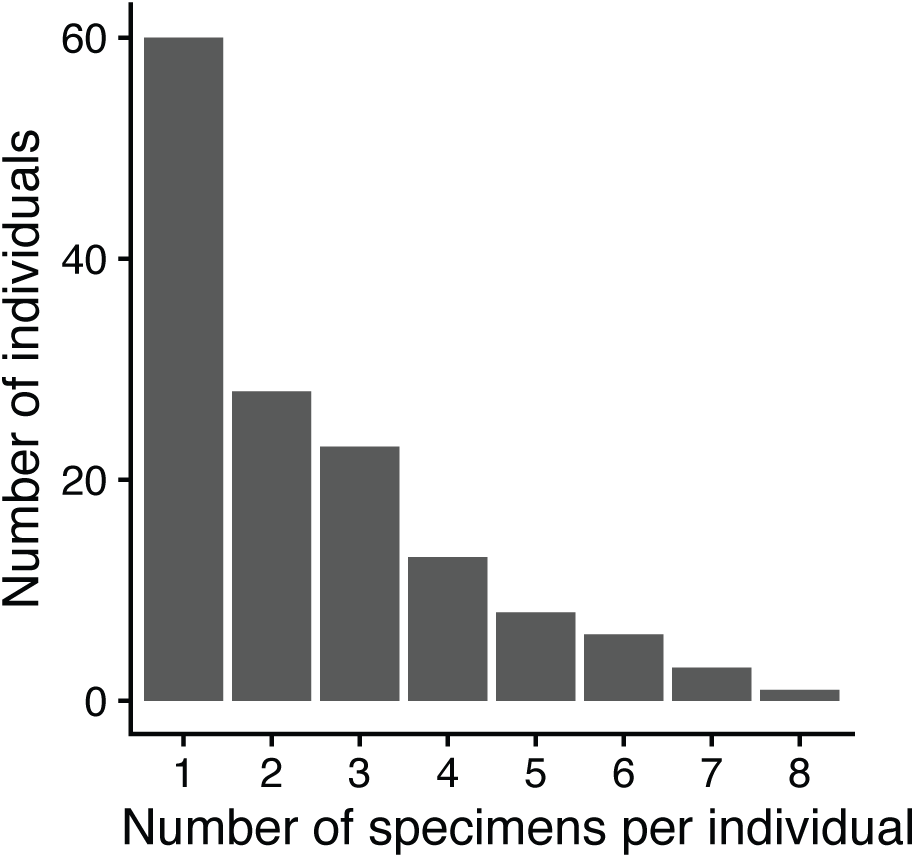
The number of specimens per individual that were successfully sequenced.

**Figure S2.**
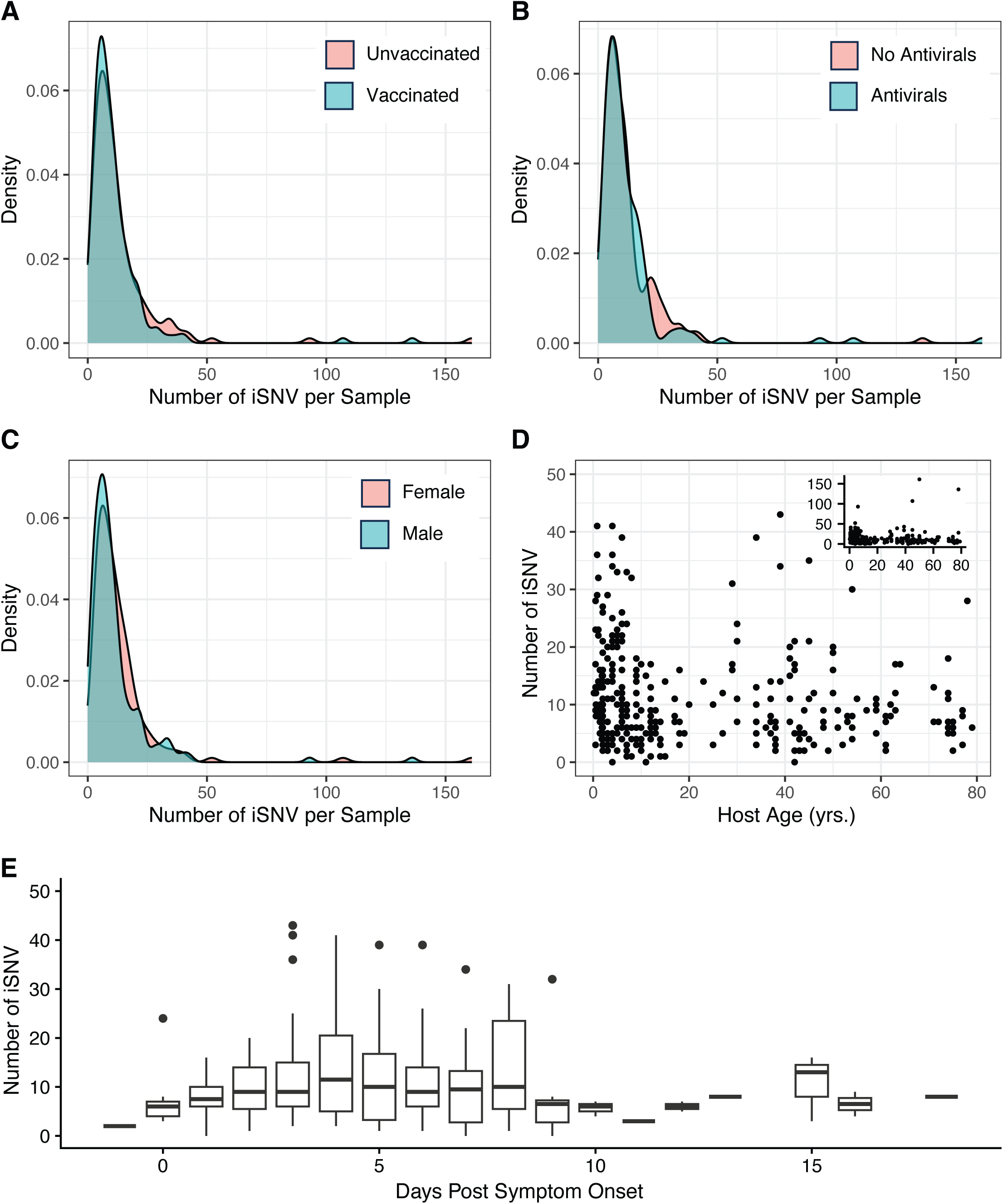
The distribution of iSNV per specimen doesn’t differ by host vaccination **(A)**, antiviral usage **(B)**, and sex **(C). (D).** The Number of iSNV per specimen by host age. The main graph is up to 50 iSNV per specimen, and the inset shows the whole range of iSNV per specimen. (**E)** The number of iSNV per specimen by day post symptom onset. For visualization purposes, specimens with greater than 50 iSNV are not shown.

**Figure S3.**
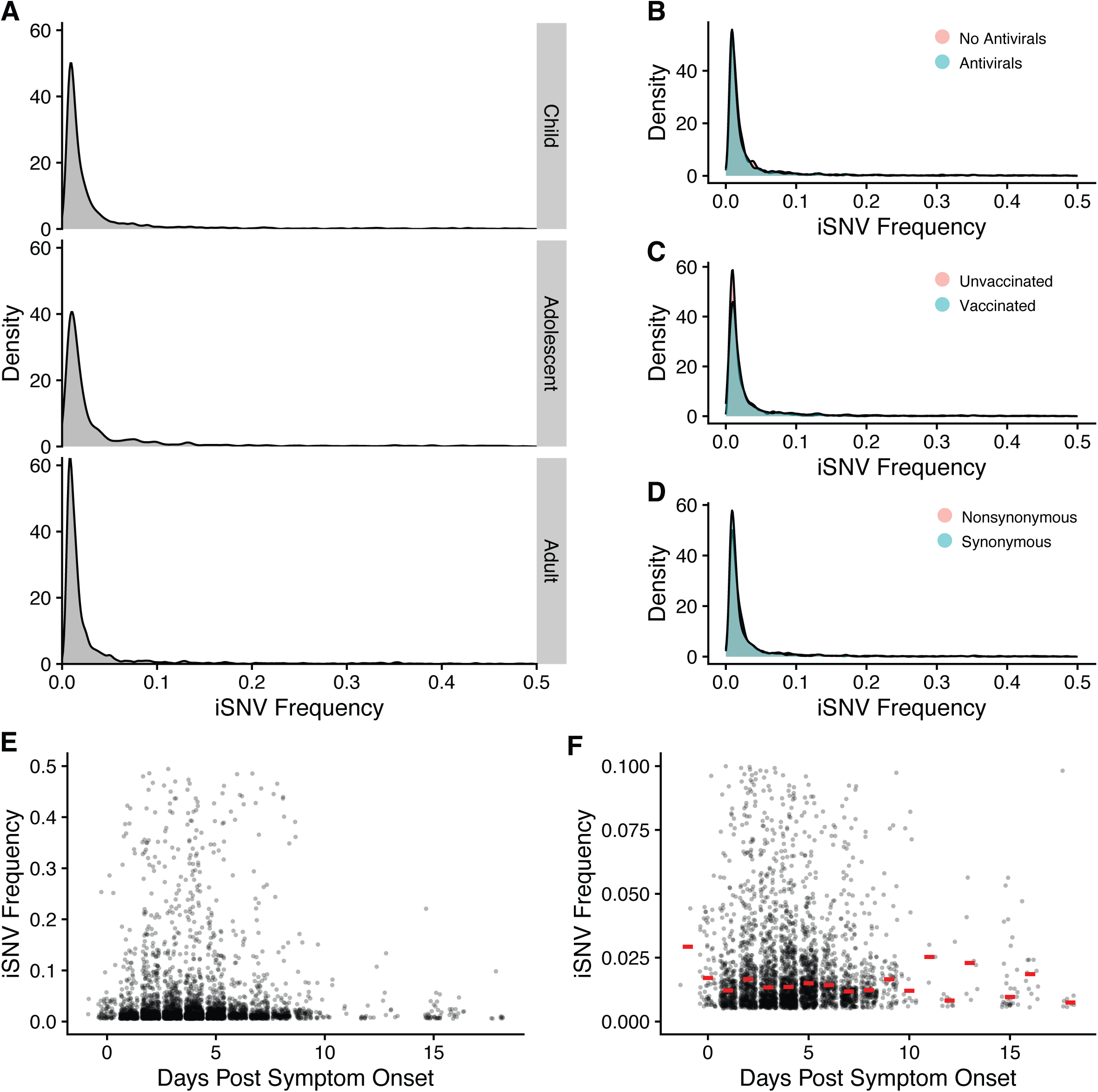
Frequency distributions of iSNV. **(A)** iSNV frequency by host age. Young children are ≤ 5 and under, adolescents are 6-18, adults are >18. iSNV frequency by antiviral usage **(B)**, vaccine status **(C),** and mutation type **(D)**. **(E).** iSNV frequency by day post symptom onset. **(F)** frequencies between 0.005 and 0.1. The red bars are the median frequency.

**Figure S4.**
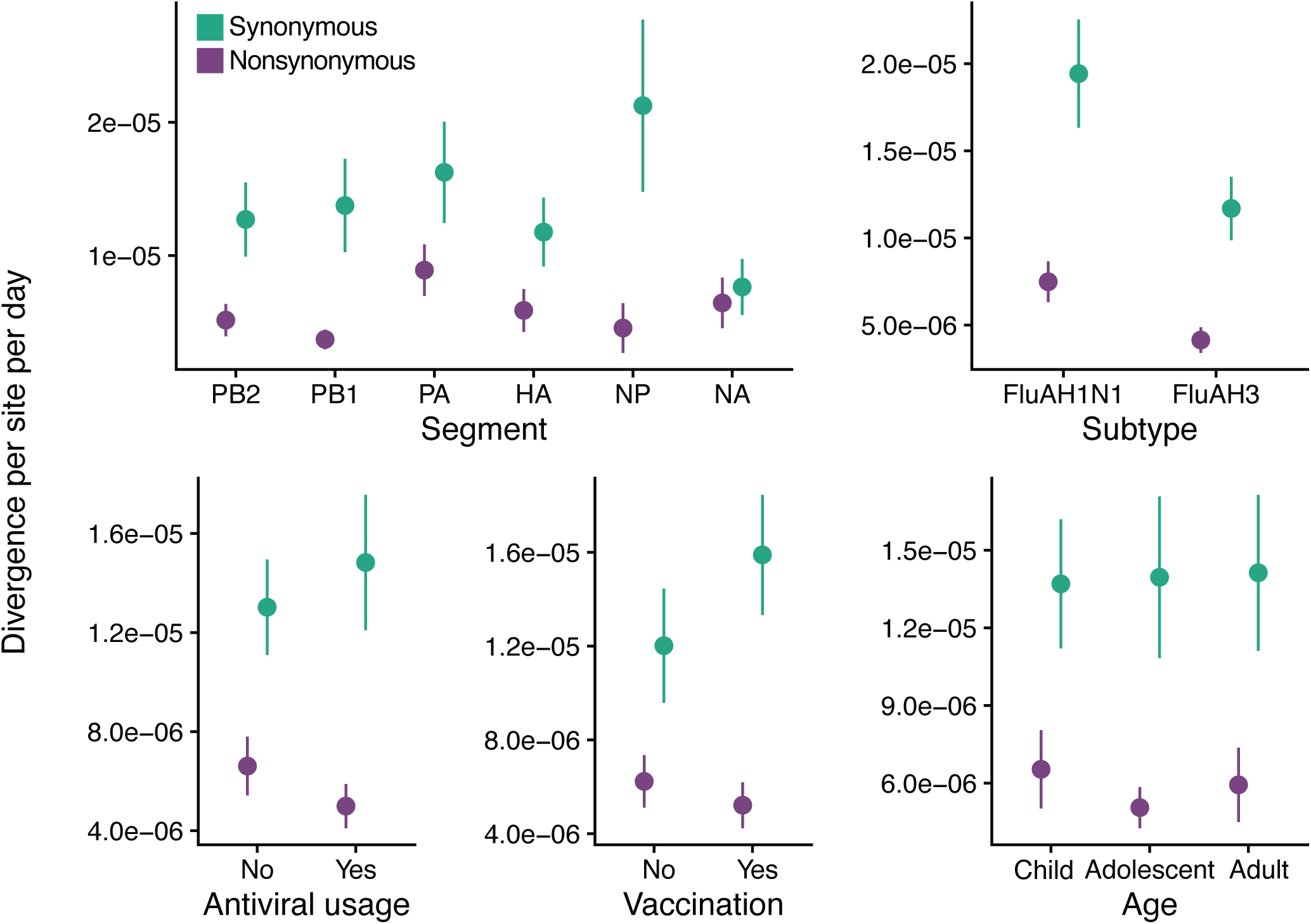
The average divergence rate by segment, subtype, host antiviral usage, age, and vaccination. All points are the mean +/- the standard error. (Green synonymous iSNV, purple nonsynonymous)

**Figure S5.**
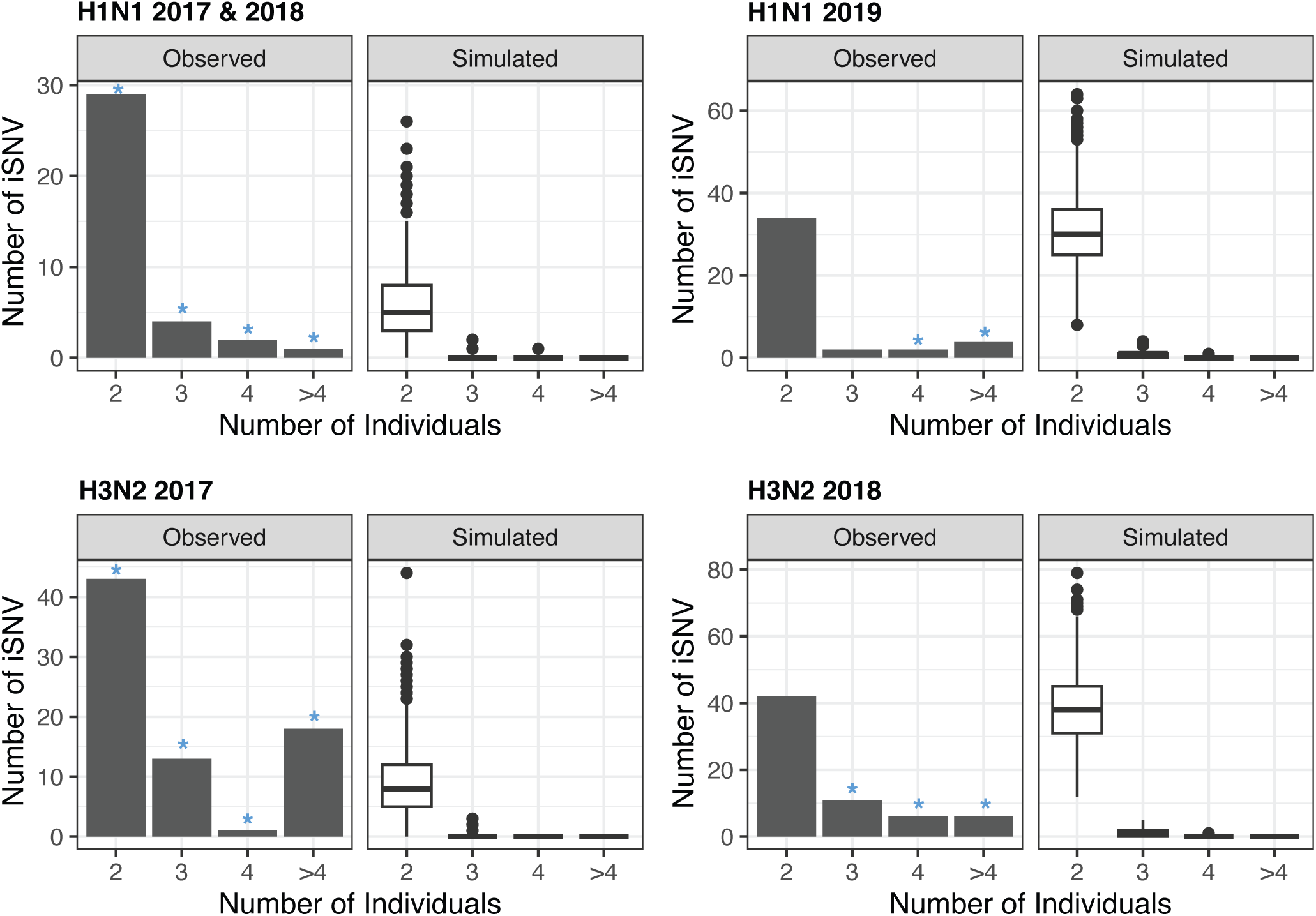
The number of iSNV that were shared across clinical specimens in the indicated number of individuals compared to simulated data (see methods). Blue asterisks indicate significant (p < 0.001) differences between the observed and simulated.

**Figure S6.**
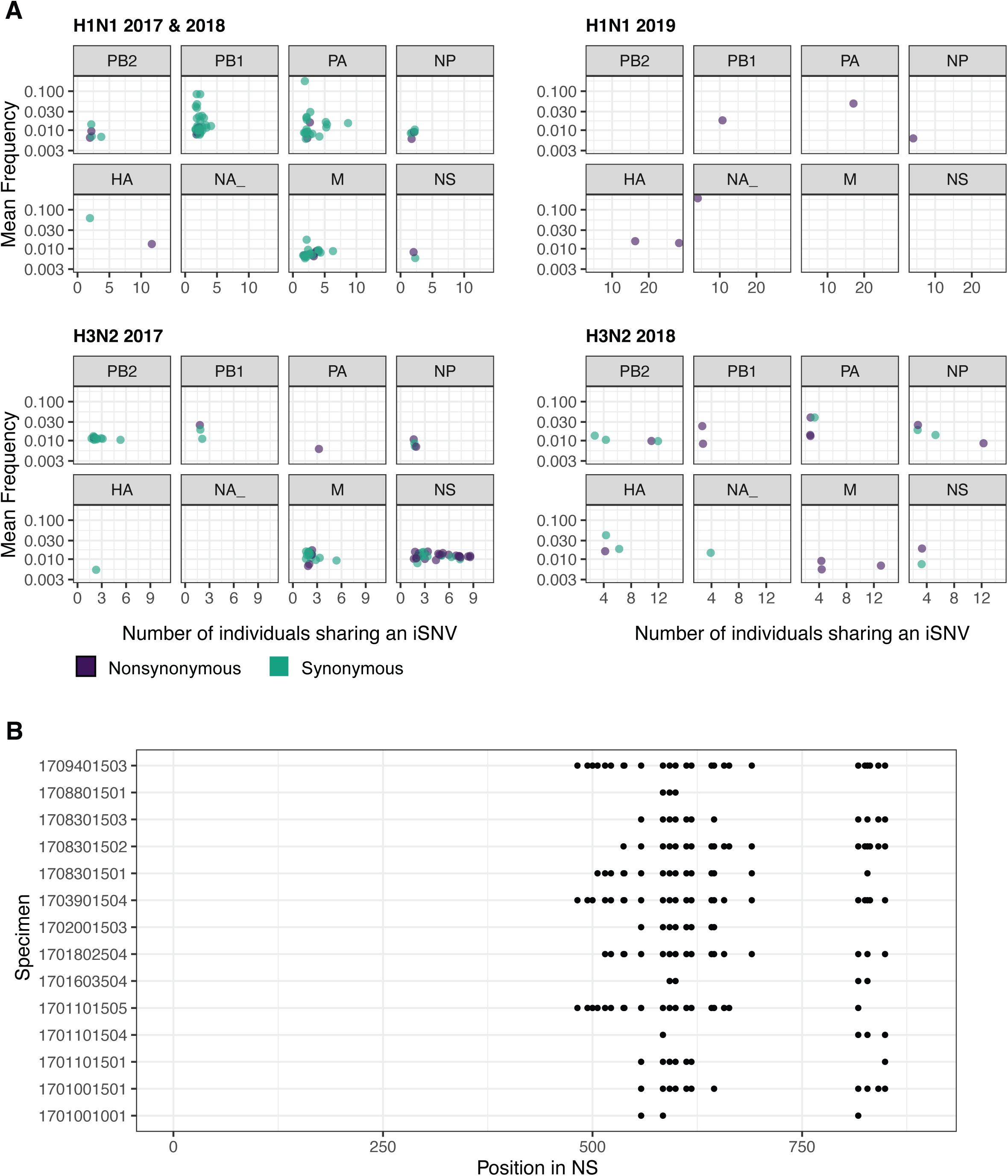
**(A).** The mean frequency of iSNV that were found in multiple individuals from different households. iSNV from 2017 & 2018 H1N1 and 2017 H3N2 infections are shared in ≥ 2 individuals. iSNV from 2019 H1N1 infections are shared in ≥ 3 individuals, and iSNV from 2018 H3N2 infections are shared in ≥ 4 individuals. (Green synonymous iSNV, purple nonsynonymous). **(B).** Position of shared iSNV (≥ 2) by sample for iSNV in NS from H3N2 2017 infections after samples with >50 iSNV were removed.

**Figure S7.**
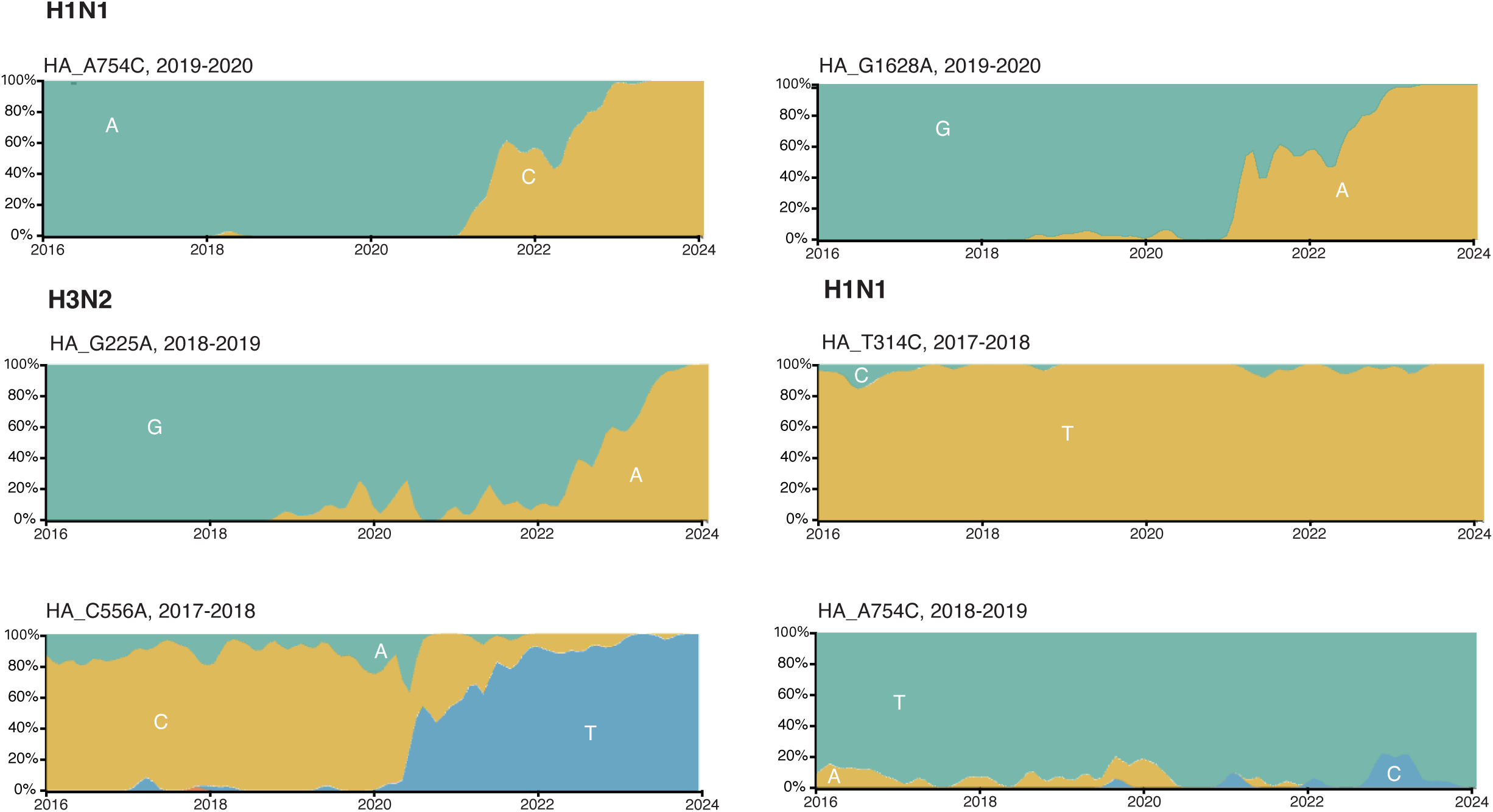
Global frequencies of HA alleles identified as under positive selection.

**Table S1.** Comparisons of the number of iSNV per specimen and of iSNV frequency. For statistically significant differences the p values are bolded. χ^2^ test statistics are from Kruskal-Wallis rank sum tests and W test statistics are from Mann-Whitney U tests.

| **Number of iSNV per sample** | Test statistic ($\chi^2$ or W) | df | p value |
| --- | --- | --- | --- |
| Age Class | 13.285 | 2 | **0.001** |
| Vaccination | 15110 | 1 | 0.228 |
| Antiviral Usage | 15158 | 1 | 0.555 |
| Days Post Symptom Onset | 17.556 | 17 | 0.417 |
| Mutation Type | 61023 | 1 | **<0.001** |
| ***Age Class - Post Hoc*** | Test statistic (Z) | p.unadj | p.adj |
| Child vs Adolescent | 3.627455 | **<0.001** | **<0.001** |
| Child vs Adult | 2.150786 | 0.031 | 0.063 |
| Adolescent vs Adult | -1.538681 | 0.124 | 0.124 |
| **iSNV Frequency** | Test statistic ($\chi^2$ or W) | df | p value |
| Age Class | 43.417 | 2 | **<0.001** |
| Vaccination | 1887183 | 1 | **< 0.001** |
| Antiviral Usage | 2225826 | 1 | **0.013** |
| Days Post Symptom Onset | 62.392 | 17 | **< 0.001** |
| Mutation Type | 2308742 | 1 | **0.005** |
| ***Age Class - Post Hoc*** | Test statistic (Z) | p.unadj | p.adj |
| Child vs Adolescent | -2.152914 | 0.031 | **0.031** |
| Child vs Adult | 4.494324 | < 0.001 | **< 0.001** |
| Adolescent vs Adult | 6.305439 | < 0.001 | **< 0.001** |

**Table S2.** Comparisons of divergence rates. Statistically significant differences are bolded. χ^2^ test statistics are from Kruskal-Wallis rank sum tests and W test statistics are from Mann-Whitney U tests.

|  | Nonsynonymous |  |  | Synonymous |  |  |
| --- | --- | --- | --- | --- | --- | --- |
| | Test statistic<br>($\chi^2$ or W) | df | p value | Test statistic<br>($\chi^2$ or W) | df | p value |
| Segment | 6.4119 | 5 | 0.268 | 3.4014 | 5 | 0.638 |
| Subtype | 2163 | 1 | **0.011** | 1745 | 1 | 0.078 |
| Age Class | 11.635 | 2 | **0.003** | 8.0787 | 2 | **0.018** |
| Days Post Symptom Onset | 31.413 | 17 | **0.018** | 41.389 | 17 | **0.008** |
| Vaccination | 1648 | 1 | 0.700 | 1629.5 | 1 | 0.609 |
| Antiviral Usage | 2063 | 1 | 0.080 | 2072 | 1 | 0.065 |
| **Age Class - Post Hoc** | Test statistic (Z) | p.unadj | p.adj | Test statistic (Z) | p.unadj | p.adj |
| Child vs Adolescent | 3.403515 | **< 0.001** | **0.002** | 2.841021 | 0.005 | **0.013** |
| Child vs Adult | 1.741911 | 0.081 | 0.081 | 1.680322 | 0.093 | 0.186 |
| Adolescent vs Adult | -1.744294 | 0.081 | 0.162 | -1.211188 | 0.226 | 0.226 |

**Table S3.** Chi Squared tests for the probability of having a shared variant. Statistically significant comparisons are bolded.

|  | chi-squared | df | p value |
| --- | --- | --- | --- |
| Age Class | 9.5451 | 2 | **0.008** |
| Vaccination | 2.6569 | 1 | 0.103 |
| Antiviral Usage | 1.3851 | 1 | 0.172 |
| ***Age Class - Post Hoc*** | <u>chi-squared</u> | <u>p.unadj</u> | <u>p.adj</u> |
| Child vs Adolescent | 1.1357 | 0.287 | 0.859 |
| Child vs Adult | 9.3301 | 0.002 | 0.068 |
| Adolescent vs Adult | 2.5414 | 0.111 | 0.333 |

**Table S4.** Chi Squared tests for the probability of an individual having at least 1 iSNV under positive selection.

|  | chi-squared | df | p-value |
| --- | --- | --- | --- |
| Age Class | 0.7067 | 2 | 0.702 |
| Vaccination | 0.367 | 1 | 0.554 |
| Antiviral Usage | 0.3329 | 1 | 0.564 |

## Citations

1. Smith, D. J. et al. Mapping the antigenic and genetic evolution of influenza virus. Science 305, 371–376 (2004).

2. Bedford, T. et al. Global circulation patterns of seasonal influenza viruses vary with antigenic drift. Nature 523, 217–220 (2015).

3. De, P., Farley, A., Lindson, N. & Aveyard, P. Systematic review and meta-analysis: influence of smoking cessation on incidence of pneumonia in HIV. BMC Med. 11, 15 (2013).

4. Salk, J. E. & Suriano, P. C. Importance of antigenic composition of influenza virus vaccine in protecting against the natural disease; observations during the winter of 1947-1948. Am. J. Public Health Nations Health 39, 345–355 (1949).

5. Kilbourne, E. D. et al. The total influenza vaccine failure of 1947 revisited: major intrasubtypic antigenic change can explain failure of vaccine in a post-World War II epidemic. Proc. Natl. Acad. Sci. U. S. A. 99, 10748–10752 (2002).

6. Luo, S., Reed, M., Mattingly, J. C. & Koelle, K. The impact of host immune status on the within-host and population dynamics of antigenic immune escape. J. R. Soc. Interface 9, 2603–2613 (2012).

7. Volkov, I., Pepin, K. M., Lloyd-Smith, J. O., Banavar, J. R. & Grenfell, B. T. Synthesizing within-host and population-level selective pressures on viral populations: the impact of adaptive immunity on viral immune escape. J. R. Soc. Interface 7, 1311–1318 (2010).

8. McCrone, J. T. et al. Stochastic processes constrain the within and between host evolution of influenza virus. eLife 7, e35962 (2018).

9. Dinis, J. M. et al. Deep Sequencing Reveals Potential Antigenic Variants at Low Frequencies in Influenza A Virus-Infected Humans. J. Virol. 90, 3355–3365 (2016).

10. Debbink, K. et al. Vaccination has minimal impact on the intrahost diversity of H3N2 influenza viruses. PLoS Pathog. 13, e1006194 (2017).

11. Han, A. X. et al. Within-host evolutionary dynamics of seasonal and pandemic human influenza A viruses in young children. eLife 10, e68917 (2021).

12. Koel, B. F. et al. Longitudinal sampling is required to maximize detection of intrahost A/H3N2 virus variants. Virus Evol. 6, veaa088 (2020).

13. Xue, K. S. & Bloom, J. D. Linking influenza virus evolution within and between human hosts. Virus Evol. 6, veaa010 (2020).

14. Sobel Leonard, A., et al. Deep Sequencing of Influenza A Virus from a Human Challenge Study Reveals a Selective Bottleneck and Only Limited Intrahost Genetic Diversification. J. Virol. 90, 11247–11258 (2016).

15. Ng, S. et al. The Timeline of Influenza Virus Shedding in Children and Adults in a Household Transmission Study of Influenza in Managua, Nicaragua. Pediatr. Infect. Dis. J. 35, 583–586 (2016).

16. Bodewes, R. et al. Prevalence of antibodies against seasonal influenza A and B viruses in children in Netherlands. Clin. Vaccine Immunol. CVI 18, 469–476 (2011).

17. Foll, M. et al. Influenza virus drug resistance: a time-sampled population genetics perspective. PLoS Genet. 10, e1004185 (2014).

18. Sobel Leonard, A., Weissman, D. B., Greenbaum, B., Ghedin, E. & Koelle, K. Transmission Bottleneck Size Estimation from Pathogen Deep-Sequencing Data, with an Application to Human Influenza A Virus. J. Virol. 91, e00171–17 (2017).

19. Doud, M. B., Hensley, S. E. & Bloom, J. D. Complete mapping of viral escape from neutralizing antibodies. PLoS Pathog. 13, e1006271 (2017).

20. Rolfes, M. A. et al. Household Transmission of Influenza A Viruses in 2021-2022. JAMA 329, 482–489 (2023).

21. McCrone, J. T. & Lauring, A. S. Measurements of Intrahost Viral Diversity Are Extremely Sensitive to Systematic Errors in Variant Calling. J. Virol. 90, 6884–6895 (2016).

22. Langmead, B. & Salzberg, S. L. Fast gapped-read alignment with Bowtie 2. Nat. Methods 9, 357–359 (2012).

23. Picard toolkit. Broad Inst. GitHub Repos. (2019).

24. Grubaugh, N. D. et al. An amplicon-based sequencing framework for accurately measuring intrahost virus diversity using PrimalSeq and iVar. Genome Biol. 20, 8 (2019).

25. Hoffmann, E., Stech, J., Guan, Y., Webster, R. G. & Perez, D. R. Universal primer set for the full-length amplification of all influenza A viruses. Arch. Virol. 146, 2275–2289 (2001).

26. Baccam, P., Beauchemin, C., Macken, C. A., Hayden, F. G. & Perelson, A. S. Kinetics of influenza A virus infection in humans. J. Virol. 80, 7590–7599 (2006).

27. Beauchemin, C. A. A. & Handel, A. A review of mathematical models of influenza A infections within a host or cell culture: lessons learned and challenges ahead. BMC Public Health 11 Suppl 1, S7 (2011).

28. Carrat, F. et al. Time lines of infection and disease in human influenza: a review of volunteer challenge studies. Am. J. Epidemiol. 167, 775–785 (2008).

29. Valesano, A. L. et al. Influenza B Viruses Exhibit Lower Within-Host Diversity than Influenza A Viruses in Human Hosts. J. Virol. 94, e01710–19 (2020).

30. Visher, E., Whitefield, S. E., McCrone, J. T., Fitzsimmons, W. & Lauring, A. S. The Mutational Robustness of Influenza A Virus. PLoS Pathog. 12, e1005856 (2016).

31. Foll, M., Shim, H. & Jensen, J. D. WFABC: a Wright-Fisher ABC-based approach for inferring effective population sizes and selection coefficients from time-sampled data. Mol. Ecol. Resour. 15, 87–98 (2015).

32. Smith, B. J. boa: An R Package for MCMC Output Convergence Assessment and Posterior Inference. J. Stat. Softw. 21, 1–37 (2007).

33. Garten, R. et al. Update: Influenza Activity in the United States During the 2017-18 Season and Composition of the 2018-19 Influenza Vaccine. MMWR Morb. Mortal. Wkly. Rep. 67, 634–642 (2018).

34. Xu, X. et al. Update: Influenza Activity in the United States During the 2018-19 Season and Composition of the 2019-20 Influenza Vaccine. MMWR Morb. Mortal. Wkly. Rep. 68, 544– 551 (2019).

35. Dawood, F. S. et al. Interim Estimates of 2019-20 Seasonal Influenza Vaccine Effectiveness - United States, February 2020. MMWR Morb. Mortal. Wkly. Rep. 69, 177–182 (2020).

36. Sobel Leonard, A., et al. The effective rate of influenza reassortment is limited during human infection. PLoS Pathog. 13, e1006203 (2017).

37. Hadfield, J. et al. Nextstrain: real-time tracking of pathogen evolution. Bioinforma. Oxf. Engl. 34, 4121–4123 (2018).

38. Morris, D. H. et al. Asynchrony between virus diversity and antibody selection limits influenza virus evolution. eLife 9, e62105 (2020).

39. Xue, K. S. et al. Parallel evolution of influenza across multiple spatiotemporal scales. eLife 6, e26875 (2017).

40. Kiso, M. et al. Resistant influenza A viruses in children treated with oseltamivir: descriptive study. Lancet Lond. Engl. 364, 759–765 (2004).

41. Gubareva, L. V., Kaiser, L., Matrosovich, M. N., Soo-Hoo, Y. & Hayden, F. G. Selection of influenza virus mutants in experimentally infected volunteers treated with oseltamivir. J. Infect. Dis. 183, 523–531 (2001).

42. FIsher, R. The Genetical Theory Of Natural Selection. (At The Clarendon Press, 1930).

43. Worby, C. J. et al. On the relative role of different age groups in influenza epidemics. Epidemics 13, 10–16 (2015).

44. Varble, A. et al. Influenza A Virus Transmission Bottlenecks Are Defined by Infection Route and Recipient Host. Cell Host Microbe 16, 691–700 (2014).

45. Lakdawala, S. S. et al. The soft palate is an important site of adaptation for transmissible influenza viruses. Nature 526, 122–125 (2015).

46. Richard, M. et al. Influenza A viruses are transmitted via the air from the nasal respiratory epithelium of ferrets. Nat. Commun. 11, 766 (2020).

47. Amato, K. A. et al. Influenza A virus undergoes compartmentalized replication in vivo dominated by stochastic bottlenecks. Nat. Commun. 13, 3416 (2022).

48. Hammond, A. et al. Review of the 2018–2019 influenza season in the northern hemisphere. Wkly. Epidemiol. Rec. 32, 345–363 (2019).

49. Hammond, A. et al. Review of the 2017–2018 influenza season in the northern hemisphere. Wkly. Epidemiol. Rec. 34, 429–444 (2018).

50. Karlsson, E. A. et al. Review of global influenza circulation, late 2019 to 2020, and the impact of the COVID-19 pandemic on influenza circulation. Wkly. Epidemiol. Rec. 25, 241–264 (2021).

51. Dhanasekaran, V. et al. Human seasonal influenza under COVID-19 and the potential consequences of influenza lineage elimination. Nat. Commun. 13, 1721 (2022).

52. Braun, K. M. et al. Acute SARS-CoV-2 infections harbor limited within-host diversity and transmit via tight transmission bottlenecks. PLOS Pathog. 17, e1009849 (2021).

53. Hannon, W. W. et al. Narrow transmission bottlenecks and limited within-host viral diversity during a SARS-CoV-2 outbreak on a fishing boat. Virus Evol. 8, 1–9 (2022).

54. Bendall, E. E. et al. Rapid transmission and tight bottlenecks constrain the evolution of highly transmissible SARS-CoV-2 variants. Nat. Commun. 14, 272 (2023).

55. Lythgoe, K. A. et al. SARS-CoV-2 within-host diversity and transmission. Science 372, eabg0821 (2021).

56. Farjo, M. et al. Within-host evolutionary dynamics and tissue compartmentalization during acute SARS-CoV-2 infection. J. Virol. 98, e0161823 (2024).

57. Lin, G.-L. et al. Distinct patterns of within-host virus populations between two subgroups of human respiratory syncytial virus. Nat. Commun. 12, 5125 (2021).

